# Predicting cellular aging following exposure to adversity: Does accumulation, recency, or developmental timing of exposure matter?

**DOI:** 10.1101/355743

**Authors:** Sandro Marini, Kathryn A. Davis, Thomas W. Soare, Matthew J. Suderman, Andrew J. Simpkin, Andrew D.A.C. Smith, Erika J. Wolf, Caroline L. Relton, Erin C. Dunn

## Abstract

Exposure to adversity has been linked to accelerated biological aging, which in turn has been shown to predict numerous health problems, including neuropsychiatric disease. In recent years, measures of DNA methylation-based epigenetic age – known as “epigenetic clocks” – have been used to estimate accelerated epigenetic aging. Yet, few studies have been conducted in children. Using data from the Avon Longitudinal Study of Parents and Children (n=973), we explored the prospective association between repeated measures of childhood exposure to seven types of adversity on epigenetic age assessed at age 7 using the Horvath and Hannum epigenetic clocks. With a Least Angle Regression variable selection procedure, we evaluated the effects of the developmental timing, accumulation, and recency of adversity exposure. We found that exposure to sexual or physical abuse, financial stress, or neighborhood disadvantage during sensitive periods in early and middle childhood best explained variability in the deviation of the Hannum epigenetic age from the chronological age. Secondary sex-stratified analyses identified particularly strong sensitive period effects, such that by age 7, girls who were exposed to abuse at age 3.5 were biologically older than their unexposed peers by almost 2 months. These effects were undetected in analyses comparing children “exposed” versus “unexposed” to adversity. Our results suggest that exposure to adversity may alter methylation processes in ways that perturb normal cellular aging and that these effects may be heightened during sensitive periods in development. Research is needed to demonstrate the effect of accelerated epigenetic aging on negative health outcomes following childhood adversity exposure.

## Introduction

Exposure to childhood adversity, such as abuse or poverty, represents one of the most potent risk factors for a range of negative health outcomes––including mental health problems–– across the lifespan, with estimates linking such exposures to at least a two-fold increase in subsequent risk for mental disorders^1–6^. Although these associations are well-established, the specific mechanisms through which adversity “gets under the skin” remain poorly understood.

Accumulating evidence suggests adversity may become biologically embedded through accelerated aging of cells, tissues, and organs^7–9^. Accelerated biological aging, in which biological age outpaces chronological age, is known to be a valid indicator of both the impaired functionality of the cell, and the biological system in which the cell interacts^10^. To date, telomere length has been the most studied measure of biological aging^11,12^, with dozens of studies showing an association between psychosocial stressors – ranging from maternal depression^13^to child maltreatment^14^to neighborhood disorder^15^– and shorter telomere length in children^16^and adults^17^. One meta-analysis of this association in adults found that a broadly defined self-reported measure of psychological stress was moderately and negatively (r=−0.25) correlated with telomere length^18^.

More recently, DNA methylation (DNAm) patterns at specific CpG sites have been proposed as a promising alternative measure of biological aging. These DNAm-based measures are referred to as the “epigenetic clocks” due to their remarkably high correlation with chronological age (r =0.96) ^19,20^. Two independent algorithms that have been developed to generate these DNAm-based age estimates are: the Horvath clock^19^, which uses multi-tissue DNAm signatures at 353 loci, and the Hannum clock^20^, which uses whole-blood DNAm measures at 71 loci located in or near genes associated with oxidative stress, DNA damage and repair, and/or the development of age-related disease. Both clocks can be used to capture accelerated epigenetic aging, which represents the discrepancy between the estimate of epigenetic age based on DNAm patterns and an individual’s actual chronological age^19,20^. Although both epigenetic clocks were first developed in adults, studies have also confirmed their appropriateness in younger populations^21,22^. Accelerated epigenetic aging as measured by these epigenetic clocks has been correlated with numerous adverse health outcomes, including frailty^23^, and increased mortality risk, such that individuals with accelerated aging show around a 20% higher mortality risk over 10 years in comparison to those without epigenetic age acceleration^24,25^.

Yet, despite promising evidence of the predictive power of epigenetic clocks and the association between adversity exposure and other forms of biological aging, only a handful of studies to date have explored how exposure to adversity influences epigenetic aging. One study of children between 6-13 years of age found that children who had twice as much exposure to violence had older epigenetic age profiles than their less exposed peers^26^. Studies of adults have also found links between cumulative lifetime stress^7^, childhood exposure to parental depression^27,28^, sexual abuse^29^, and chronic financial stress^30^with accelerated epigenetic aging.

Although evidence from these studies suggests a link between exposure to adversity and epigenetic aging, most of this work has primarily focused on one or two types of adversity, as opposed to a range of possible exposure types. Furthermore, to our knowledge, no studies have examined the importance of the characteristics of adversity, including the timing and duration of exposure. Given the growing body of support for “sensitive periods” in development, during which time developing organs, tissues, and biological systems may be particularly susceptible to the effects of experience^31–33^, as well as evidence documenting the consequences of more accumulated exposure^34^, consideration of the timing and duration of adversity across the life course is warranted.

In the current study, we aimed to address these limitations by: (1) investigating the association between exposure to multiple types of adversity between birth and age 7 on epigenetic aging at age 7, and (2) identifying the life course theoretical models that best explained the relationship between adversity exposure and epigenetic aging, for each adversity type. To accomplish these goals, we used data from a prospective, longitudinal study of children to test three theoretical models derived from life course theory^35,36^: a *sensitive or critical period model*, which posits that the developmental timing of exposure is most important in shaping accelerated aging^32,37^, an *accumulation model*, which posits that every additional year of exposure is associated with an increased risk for accelerated aging^34,38,39^, and a *recency* model, which suggests that the effects of adversity can be time-limited, and thus accelerated epigenetic aging may be more strongly linked to proximal rather than distal events^40^. Three secondary analyses focused on evaluating the importance of studying these adversity characteristics relative to simply examining the presence or absence of exposure, using a broader set of age ranges to define sensitive periods, and understanding sex-specific effects.

## Methods

### Sample and Procedures

We used data from the Avon Longitudinal Study of Parents and Children (ALSPAC), a large population-based birth cohort out of Avon, England of children followed from before birth through early adulthood^41–43^. ALSPAC generated blood-based DNAm profiles at age 7 as part of the Accessible Resource for Integrated Epigenomics Studies (ARIES), which is a subsample of 1,018 mother-child pairs from the ALSPAC^44^. These ARIES mother-child pairs were randomly selected out of those with complete data across at least five waves of data collection. See **Supplemental Materials** for additional information about the ALSPAC sample.

### Measures

#### Cellular Aging

DNAm was determined at age 7 using the Illumina Human Methylation 450k BeadChip microarray, which captures DNAm at 99% of RefSeq genes (over 485,000 CpG sites). DNA methylation wet laboratory procedures, preprocessing analyses, and quality control were performed at the University of Bristol as described elsewhere^44^. The level of methylation is expressed as a ‘beta’ value (β-value), representing the proportion of cells methylated at each interrogated CpG site, and ranges from 0 (no methylated dinucleotides observed) to 1 (all dinucleotides methylated).

Using the β-values for each participant in the sample, we generated two estimates of epigenetic age based on the approaches of Horvath^19^and Hannum^20^. For each clock, we estimated age acceleration using a regression procedure in which epigenetic age was the outcome and chronological age was the independent variable. For the Horvath clock, we used the online epigenetic clock calculator (http://labs.genetics.ucla.edu/horvath/dnamage/) to calculate “intrinsic epigenetic age acceleration” (IEAA), which is derived by regressing the epigenetic age against chronological age, adjusting for cell counts ^45^. To derive the Hannum clock, we summed the normalized β-values using the Touleimat method^46^multiplied by the 71 respective regression coefficients obtained by Hannum and colleagues in their model^20^. This regression procedure adjusted for blood cell composition (specifically percentage of CD8+,CD4+,CD56, CD19,CD14, and granulocytes), similar to the IEAA score. In both the Horvath and Hannum epigenetic clocks, age acceleration or deceleration is represented by the residuals of the above described regression procedures ^24,47^. Positive residuals indicate accelerated aging, in which the child’s chronological age is lower than their estimated methylation age (hereafter referred to as accelerated aging). Conversely, negative residuals indicate age deceleration, in which the child’s estimated methylation age is lower than their actual chronological age.

#### Exposure to Adversity

We examined the effect of seven adversities on methylation age residuals: (a) caregiver physical or emotional abuse; (b) sexual or physical abuse (by anyone); (c) maternal psychopathology; (d) one adult in the household; (e) family instability; (f) financial stress/poverty; and (g) neighborhood disadvantage/poverty. These adversity types were chosen based on previous research^29,48–50^linking these exposures to epigenetic change^27,29,48,49^or accelerated biological aging^13–16^. Each type of adversity was measured on at least five occasions at or before age 7 (see Table 1) from a single item or psychometrically validated standardized measures. For each type of adversity, we generated three sets of variables to test the three life course hypotheses: (a) for the *sensitive period hypothesis*, we created a set of variables indicating presence versus absence of the adversity at a specific developmental stage; specific time periods of assessment for each adversity are denoted in **Supplemental Table 1.** To test the (b) *accumulation hypothesis*, we generated a single variable denoting the total number of time periods of exposure to a given type of adversity. For the (c) *recency hypothesis*, we generated a single variable denoting the total number of developmental periods of exposure, with each exposure weighted by the age in months of the child during the measurement time period; this recency variable gave a larger weight to more recent exposures, thus, allowing us to determine whether more recent exposures were more impactful.

**Table 1.** Exposure to childhood adversity in the total analytic sample and by the age at exposure (n=973)

|  |  |  |  |  |  |  |  |  |  |  |  |  |  |  |
| --- | --- | --- | --- | --- | --- | --- | --- | --- | --- | --- | --- | --- | --- | --- |
|  | Caregiver<br>physical or<br>emotional<br>abuse |  | Sexual or<br>physical<br>abuse (by<br>anyone) |  | Family instability |  | Maternal<br>psychopathology |  | Financial stress |  | One adult in the<br>household |  | Neighborhood<br>disadvantage |  |
|  | N | (%) | N | (%) | N | (%) | N | (%) | N | (%) | N | (%) | N | (%) |
| Unexposed | 822 | 84.48 | 850 | 87.36 | 499 | 51.28 | 686 | 70.5 | 669 | 68.76 | 833 | 85.61 | 829 | 85.2 |
| Exposed | 151 | 15.52 | 123 | 12.64 | 474 | 48.72 | 287 | 29.5 | 304 | 31.24 | 140 | 14.39 | 144 | 14.8 |
| *Age at Exposure* |  |  |  |  |  |  |  |  |  |  |  |  |  |  |
| Very early childhood |  |  |  |  |  |  |  |  |  |  |  |  |  |  |
| 8 months | 34 | 3.66 | --- | --- | --- | --- | 95 | 10.24 | 104 | 11.26 | 243 | 3.67 | --- | --- |
| 1.5/1.75 yrs | 38 | 4.15 | 28 | 3 | 170 | 18.22 | 89 | 9.81 | 98 | 10.73 | 288 | 4.32 | 76 | 8.44 |
| 2/2.75 yrs | 56 | 6.26 | 32 | 3.58 | 182 | 20.34 | 130 | 14.77 | 97 | 10.89 | 350 | 5.2 | 74 | 8.36 |
| Early childhood |  |  |  |  |  |  |  |  |  |  |  |  |  |  |
| 3.5 yrs | --- | --- | 36 | 3.96 | 186 | 20.46 | 114 | 12.9 | 140 | 14.46 | --- | --- | --- | --- |
| 4.0/4.75 yrs | 41 | 4.57 | 35 | 3.91 | 118 | 13.22 | --- | --- | --- | --- | 410 | 6.88 | --- | --- |
| 5/5.75 yrs | 57 | 6.36 | 24 | 2.74 | 114 | 13.01 | --- | --- | --- | --- | --- | --- | 55 | 6.21 |
| Middle childhood |  |  |  |  |  |  |  |  |  |  |  |  |  |  |
| 6/6.75 yrs | 50 | 5.66 | 23 | 2.61 | 69 | 7.82 | 130 | 14.91 | --- | --- | --- | --- | --- | --- |
| 7 yrs | --- | --- | --- | --- | --- | --- | --- | --- | 121 | 12.5 | 504 | 7.6 | 43 | 4.89 |

##### Covariates

We controlled for the following covariates, measured at child birth: child race/ethnicity; number of births in the pregnancy (pregnancy size); number of previous pregnancies; maternal marital status; highest level of maternal education; maternal age; maternal smoking during pregnancy; child birth weight; parental homeownership; and parent job status (see **Supplemental Materials** for coding).

### Analyses

We began by running univariate and bivariate analyses to examine the distribution of covariates and exposures to adversity in the total analytic sample. To reduce potential bias and minimize loss of power due to attrition, we performed multiple imputation (on missing exposures and covariates): we used logistic regression in 20 datasets with complete data on the outcome with 25 iterations, separately for each exposure, including covariates (see **Supplemental Materials**). We then used a novel two-stage structured life course modeling approach (SLCMA)^51,52^to evaluate, separately for each adversity type, which of the three life course theoretical models (sensitive period, accumulation, recency) could best explain the relationship between adversity exposure and epigenetic age. Compared to other methods, such as standard multiple regression, the SLCMA provides an unbiased way to compare multiple competing theoretical models simultaneously and identify the most parsimonious explanation for variation in epigenetic age.

In the first stage of the SLCMA, we entered the set of variables described earlier into the Least Angle Regression (LARS) variable selection procedure^53^. LARS identifies the smallest combination of variables that explain the most amount of outcome variation. We used a covariance test^54^and elbow plots (**Supplemental Figure 1**) to determine whether the selected models were supported by the ARIES data. In the second stage, the life course theoretical models found in the first stage to best fit the observed data – that is, the model(s) appearing at the “elbow” of the plot and/or those with p-values <.05 in the covariance test – were then carried forward to a multivariate regression framework. In this framework, measures of effect were estimated for all selected hypotheses (see **Supplemental Materials** for details on LARS selection procedure). With respect to multiple testing, the covariance test p-values are adjusted for the number of variables included in the LARS procedure, controlling the type I error rate for each adversity type on each epigenetic clock.

#### Secondary Analyses

Three additional sets of analyses were performed following the primary analyses described above. First, to gauge the importance of studying these adversity characteristics relative to simply examining the presence or absence of exposure, we compared the results derived from the three life course models to those obtained from an ever versus never exposed model. (Table 3).

Second, to explore the possibility that a broader definition of sensitive periods would yield comparable results, and to facilitate interpretation of our findings in comparison to prior studies^55–58^, we re-analyzed our data focusing on three sensitive periods: *very early childhood*(ages 8 months – 2.75 years); *early childhood*(ages 3.5 – 5.75 years); and *middle childhood* (ages 6 – 7 years). Third, we performed sex-stratified secondary analyses, given that adversity exposure^59^varies between males and females and males overall have higher IEAA than females^22^.

## Results

### Sample Characteristics

There were 973 children in the analytic sample (50.2% female, 97.2% white). At the time of their child’s birth, most mothers were between 20-35 years of age (89.5%), married (82.6%), non-smokers (89.3%), and living in their own home (88.3%). Forty-seven percent of mothers were experiencing their first pregnancy. Descriptive statistics on other covariates are presented in **Supplemental Table 2**.

### Distribution of Exposure to Adversity and Age Acceleration

Table 1 shows the prevalence of childhood adversity overall and by each age period of assessment. The lifetime prevalence of adversity exposure ranged from 12.6% for physical abuse to 48.7% for family instability. For some adversities, including caregiver physical or emotional abuse, as well as sexual or physical abuse, the prevalence of exposure was equally distributed across the developmental time periods. For other adversities, including maternal psychopathology and financial stress, exposure was concentrated during very early childhood, meaning from 8 months to 2.75 years. Children exposed to any type of adversity were more likely than their unexposed peers to be non-white and born to non-married mothers with low education, low social class, and with more than three previous pregnancies (**Supplemental Table 2**).

As shown in **Supplemental Table 2**, girls were, on average, epigenetically older than boys. Also, children born to married mothers with higher education and lower social class had lower age residuals (according to Hannum’s epigenetic clock) compared to children whose mothers fell into other corresponding categories. No significant differences were observed for race, maternal smoking, weight at birth, maternal education, pregnancy size, home ownership, and number of previous pregnancies (all p-values >.10) (**Supplemental Table 2**).

### Association between Exposure to Adversity and Age Acceleration

Table 2 displays, separately for each adversity type and epigenetic clock, the theoretical model selected by the LARS that best explained variability in age acceleration. As shown, evidence for three associations emerged for Hannum’s epigenetic clock, all of which emphasized age acceleration following adversity exposure and the importance of sensitive periods. First, we found evidence that exposure to sexual or physical abuse at 3.5 years was associated with older epigenetic age (effect β=.07 years; 95% CI=.00-.14, p=.001, R^2^=.01). Similarly, exposure to financial stress at 7 years (effect β=.11, CI=.08-.14, p = .001, R^2^=.05), and neighborhood disadvantage at 7 years (effect β= .12 years, CI=.01-.22, p = .001, R^2^=.01) were associated with an acceleration in epigenetic aging. The magnitude of these beta estimates translates into an age acceleration of about one month among children exposed to adversity. None of the other life course theoretical models were selected as explaining a significant amount of the variability in age acceleration for these three or any other adversity types. Using Horvath’s epigenetic clock, none of the life course models were associated with epigenetic age acceleration for any of the adversities studied (Table 2).

**Table 2.**
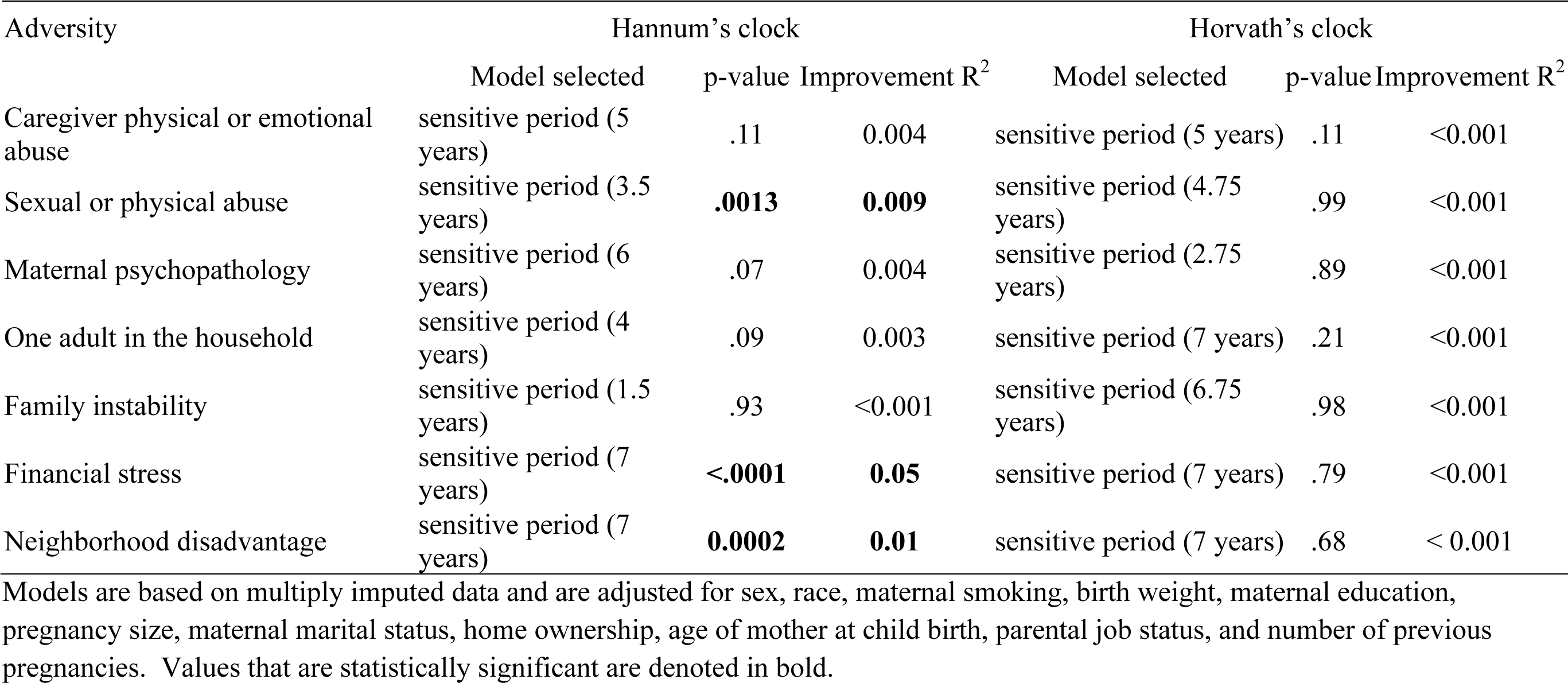
Results of LARS models showing the life course theoretical model that best explained the relationship between adversity and age acceleration (n=973)

| Adversity | Hannum's clock |  |  | Horvath's clock |  |  |
| --- | --- | --- | --- | --- | --- | --- |
|  | Model selected | p-value | Improvement R<sup>2</sup> | Model selected | p-value | Improvement R<sup>2</sup> |
| Caregiver physical or emotional abuse | sensitive period (5 years) | .11 | 0.004 | sensitive period (5 years) | .11 | <0.001 |
| Sexual or physical abuse | sensitive period (3.5 years) | **.0013** | **0.009** | sensitive period (4.75 years) | .99 | <0.001 |
| Maternal psychopathology | sensitive period (6 years) | .07 | 0.004 | sensitive period (2.75 years) | .89 | <0.001 |
| One adult in the household | sensitive period (4 years) | .09 | 0.003 | sensitive period (7 years) | .21 | <0.001 |
| Family instability | sensitive period (1.5 years) | .93 | <0.001 | sensitive period (6.75 years) | .98 | <0.001 |
| Financial stress | sensitive period (7 years) | **<.0001** | **0.05** | sensitive period (7 years) | .79 | <0.001 |
| Neighborhood disadvantage | sensitive period (7 years) | **0.0002** | **0.01** | sensitive period (7 years) | .68 | < 0.001 |

Models are based on multiply imputed data and are adjusted for sex, race, maternal smoking, birth weight, maternal education, pregnancy size, maternal marital status, home ownership, age of mother at child birth, parental job status, and number of previous pregnancies. Values that are statistically significant are denoted in bold.

### Secondary Analyses

We performed three secondary analyses of epigenetic age acceleration. First, to compare the benefits of an aforementioned complex model of adversity versus a simpler one, we conducted an *ever vs. never exposed* model for each adversity type and found that financial stress was the only adversity associated with age acceleration, as estimated by Hannum’s clock (Table 3). This finding suggests that exposure to financial stress is associated with older epigenetic age relative to chronological age, regardless of timing. Exposure to any adversity was not associated with epigenetic age acceleration when Horvath’s clock was used (**Supplemental Table 3)**.

**Table 3.** Results of linear regression analysis of exposed vs. non-exposed on Hannum’s epigenetic clock (n=973)

|  | Beta<br>(years) | se | p-value | 95% C.I. |
| --- | --- | --- | --- | --- |
| Caregiver physical or emotional abuse | 0.012 | 0.0145 | .428 | -0.018 - 0.041 |
| Sexual or physical abuse | 0.027 | 0.015 | .078 | -0.003 - 0.056 |
| Maternal psychopathology | 0.003 | 0.011 | .812 | -0.019 - 0.025 |
| One adult in the household | 0.029 | 0.017 | .092 | -0.005 - 0.063 |
| Family instability | 0.003 | 0.011 | .748 | -0.018 - 0.024 |
| Financial stress | **0.047** | **0.010** | **<.0001** | **0.027 - 0.068** |
| Neighborhood disadvantage | 0.004 | 0.019 | .822 | -0.033 - 0.041 |

Second, we re-ran the analyses using three broader categories to define sensitive periods (very early childhood, early childhood, middle childhood). Similar results were obtained when the sensitive periods were collapsed into these three categories (**Supplemental Table 4**). In addition to the models reported in Table 2, analyses using these broader age categories found evidence of stronger associations for two adversities with Hannum’s clock. Specifically, having only one adult in the household during early childhood (effect β=.06 years, CI=.02-.09, p=.002,) and being exposed to maternal psychopathology in middle childhood (effect β=.03 years, CI=.06-.02, p =.023) was associated with a modest acceleration in epigenetic age.

Third, to evaluate potential sex-differences, we performed sex-stratified analyses which did reveal differences in the association between adversity exposure and epigenetic aging between boys and girls. Sex-stratified analyses using Hannum’s clock are reported in Table 4. For girls, having only one adult in the household, or being exposed to maternal psychopathology, financial stress, or abuse of any kind were associated with increased epigenetic age; each of these associations showed sensitive period specificity. For example, by age 7, girls who were exposed to abuse at age 3.5 were biologically older than their unexposed peers by almost 2 months. In boys, a sensitive period was identified at age 7 for exposure to financial stress and neighborhood disadvantage. Sex-stratified analyses using Horvath’s epigenetic clock are reported in **Supplemental Table 5**. This analysis revealed an association in girls between caregiver physical or emotional abuse and epigenetic age acceleration and was similar to the results using Hannum’s clock.

**Table 4.**
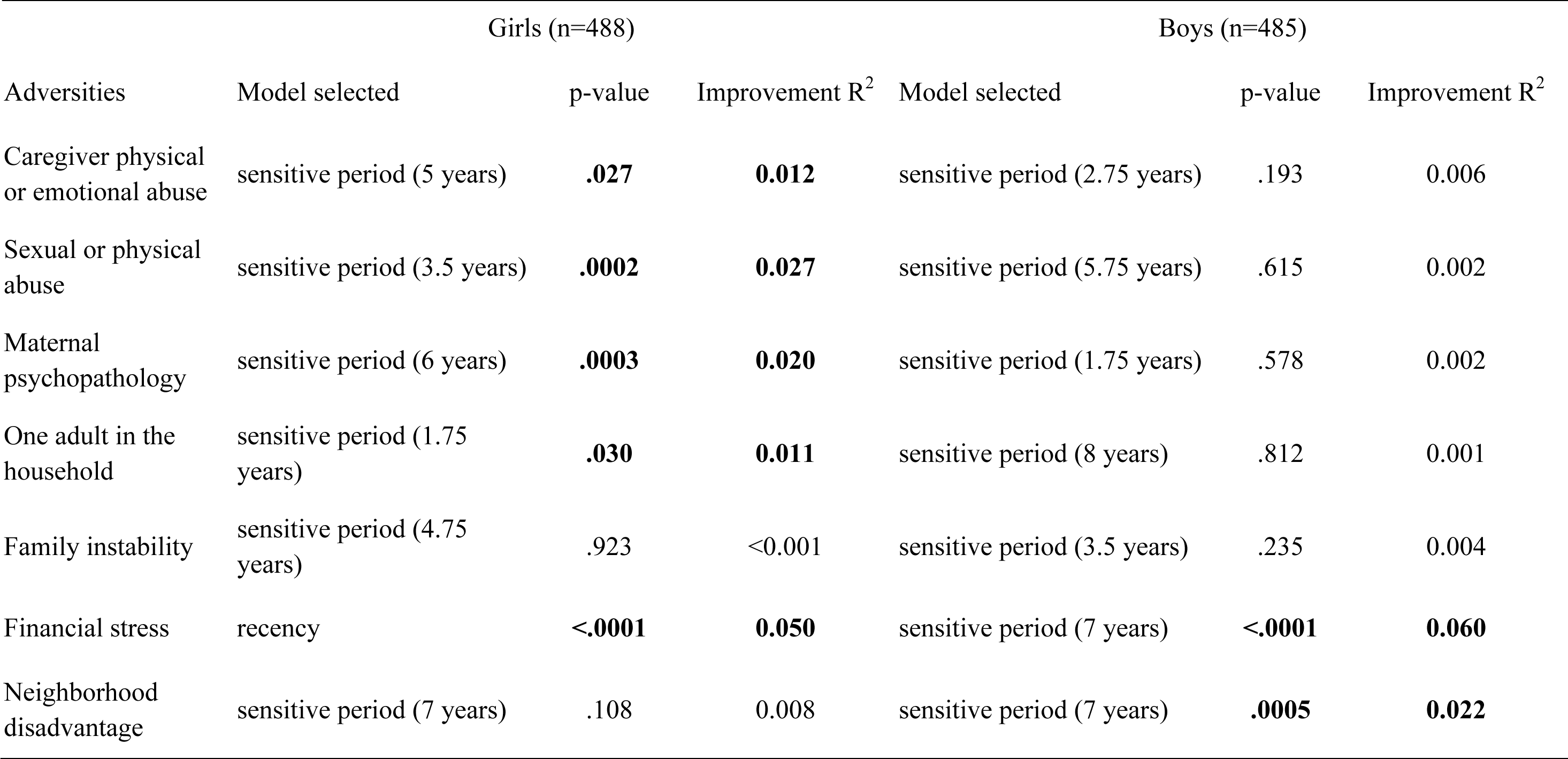
Results of LARS models showing the life course theoretical model that best explained the relationship between adversity and age acceleration, with Hannum’s epigenetic clock, stratified by sex (n=973)

| Adversities | Girls (n=488) |  |  | Boys (n=485) |  |  |
| --- | --- | --- | --- | --- | --- | --- |
|  | Model selected | p-value | Improvement R<sup>2</sup> | Model selected | p-value | Improvement R<sup>2</sup> |
| Caregiver physical or emotional abuse | sensitive period (5 years) | **.027** | **0.012** | sensitive period (2.75 years) | .193 | 0.006 |
| Sexual or physical abuse | sensitive period (3.5 years) | **.0002** | **0.027** | sensitive period (5.75 years) | .615 | 0.002 |
| Maternal psychopathology | sensitive period (6 years) | **.0003** | **0.020** | sensitive period (1.75 years) | .578 | 0.002 |
| One adult in the household | sensitive period (1.75 years) | **.030** | **0.011** | sensitive period (8 years) | .812 | 0.001 |
| Family instability | sensitive period (4.75 years) | .923 | <0.001 | sensitive period (3.5 years) | .235 | 0.004 |
| Financial stress | recency | **<.0001** | **0.050** | sensitive period (7 years) | **<.0001** | **0.060** |
| Neighborhood disadvantage | sensitive period (7 years) | .108 | 0.008 | sensitive period (7 years) | **.0005** | **0.022** |

## Discussion

This study explored the association between a variety of adversity exposures – as well as the timing and accumulation of those exposures – and accelerated epigenetic age as measured by two epigenetic clocks. By comparing different theoretical life course models of exposure, we could investigate which features of adversity exposure best captured the underlying signal. To our knowledge, this study represents one of the first to investigate whether the effects of adversity on epigenetic aging are observable in children and the extent to which these relationships may vary as a function of the timing and type of exposure.

The main finding of this study was that there appear to be sensitive periods in development when a broad range of adversity exposures are associated with an acceleration in epigenetic age. Specifically, we found that exposure to sexual or physical abuse in early childhood (age 3.5 years) and exposure to financial stress or neighborhood disadvantage in middle childhood(age 7) were all associated with epigenetic age acceleration by about one month. We acknowledge that the incremental variance explained was limited, but this estimate of effect is consistent with previous literature^60^. Although the literature to date on the association between social environmental exposures and epigenetic aging in children is limited, our findings are consistent with previous work linking adversities, such as abuse^29^, financial stress tied to chronic low income^30^, and parental psychopathology^27,29^, with accelerated epigenetic aging in adulthood.

Our results extend previous findings by exploring the effects of the timing of exposure. We found evidence for sensitive periods during early and middle childhood, when the association between adversity exposure and epigenetic aging appears to be particularly strong. This finding aligns with human^61,62^and animal^63–66^studies showing the importance of sensitive periods in epigenetic programming. It seems therefore plausible that the epigenetic age of cells is influenced by environmental inputs in a similar time-susceptibility manner. The current findings further emphasize the importance of attending to possible time-dependent effects when studying the effects of adversity on cellular aging, including DNAm and other cellular-based measures of accelerated aging. Our results suggest that an approach that does not account for the specific life stages when adversity occurs may fail to detect effects of adversity on epigenetic age acceleration, and crude classifications of children as exposed vs. unexposed to “early life” adversity may mask observed differences among those exposed to adversity.

Additionally, given known sex-differences in the effects of adversity exposure, and sex-differences intrinsic to epigenetic age acceleration, we performed a set of sex-stratified analyses. These analyses revealed that adversity could differentially affect epigenetic age acceleration in boys and girls. Specifically, we found the link between maternal psychopathology exposure and accelerated aging was only present for girls, as was the association between exposure to abuse and age acceleration. Some of these associations were particularly relevant; for example by age 7, girls who were exposed to abuse at age 3.5 were biologically older than their unexposed peers by almost 2 months. These findings suggest that the associations found in our main analyses may have been largely driven by the strength of the effect in girls. Our sex-stratified results are also consistent with previous findings indicating sex-specific effects in the patterning of epigenetic marks following childhood adversity^61,67^, and underscore the value of sex-stratification in future analyses.

It is worth noting that epigenetic age represents one of a number of biological age predictors, such as telomere length. Although more work is required, studies using both markers suggest that epigenetic and telomeric clocks may capture different but complementary aspects of biological aging^25^. For example, accelerated epigenetic aging has been correlated with a comprehensive measure of frailty, even in the absence of a correlation between telomere length and frailty^23^. In another study, telomere length and epigenetic clock estimates were each found to be independent predictors of chronological age and mortality risk, consistent with weak, nonsignificant correlations between the two biological age measures^25^. Thus, studies that investigate epigenetic markers may contribute to our understanding of biological age beyond what more traditional telomere studies can tell us.

In the current study, we did not find an association between exposure to the studied adversities on Horvath’s epigenetic clock. Other studies using both the Horvath and Hannum clocks have similarly found that associations may exist for one clock, but not for another^47^, as recently also shown in a meta-analysis ^68^. As described by others, there are a number of possibilities for such discrepancies. For instance, the Horvath and Hannum models differ in the tissue and age of subjects used to develop them, and the loci used are largely different as well, with only a few genes overlapping between the two algorithms. Therefore, the possibility exists that each clock has a level of disease specificity that is dependent on the prioritized methylation loci, making us unlikely to capturing methylation change that is more tissue-specific. Moreover, our estimate of epigenetic age using Hannum’s clock was based on directly measured cell blood count, whereas Horvath’s clock infers cell blood count from the methylation levels. Together, these differences suggest that the two clocks may be capturing slightly different aspects of biological aging, with the Horvath clock representing overall frailty in the body, whereas Hannum’s clock may be more related to immune response^60,69^. The only exposure associated with both epigenetic clocks was caregiver abuse, among girls.

There are several strengths of the current study. We included a more inclusive and detailed assessment of adversity types; most research in the field to date has focused on single types of adversity exposure, such as parental depression or low socioeconomic status only. Moreover, we also incorporated different life course theoretical models of adversity exposure, thereby allowing us to investigate which temporal features of exposure are most strongly associated with epigenetic aging. Also, although analyses on diverse racial and ethnic samples are still lacking, the current study presents findings on a cohort of children of primarily European ancestry, adding to recent work done using the ARIES subsample^70^. Finally, most studies to date have focused on older samples, often with a median chronological age above 45 years^70^, whereas the current study focused on epigenetic aging in children.

However, our study had limitations. Our findings are based on DNA extracted from blood, which may be limiting as patterns of epigenetic change following social environmental stress exposure have been found to be tissue-specific, such that the same individual may have different Horvath’s epigenetic clock estimates for different tissues ^71^. Therefore, we cannot exclude the possibility that childhood adversities affect cell methylation in a tissue-specific pattern and that blood-based measures of DNAm may not capture methylation changes of all tissues that occur following adversity. The challenge of tissue and cell-type specificity is unfortunately a limitation of all epigenome-brain research in living human subjects. Moreover, given the structure of the data and the lack of complete overlap in adversity assessment across time, we were unable to examine the adversities all together. Attending to only one adversity type at a time may lead to overestimates of the effect of a given exposure. At the same time, by examining each type of adversity individually, we were able to identify meaningful differences in the association between adversity and accelerated aging. One challenge for future analyses will be to develop new ways to examine multiple adversities simultaneously without simply summing across number of exposures^72^. Although we used multiple imputation in an effort to reduce potential bias and minimize loss of power, we cannot rule out the possibility that missing or incomplete outcome data due to attrition may have influenced our findings. Finally, since the oldest sensitive period coincides with the most recent exposure occasion for all children, it may be difficult to discern between the oldest sensitive period and recent exposure.

In conclusion, we found that adversity experiences assessed in very early, early, and middle childhood were differentially associated with accelerated epigenetic aging at age 7. These findings suggest that accelerated epigenetic aging may function as one of the mechanisms through which childhood adversity becomes biologically embedded, and that adversity exposures during sensitive periods in childhood may have a particularly strong accelerating effect on epigenetic age. Additional research is needed to further demonstrate the other aspect of the effect of accelerated cellular aging on subsequent risk for depression and other neuropsychiatric disorders. Nevertheless, understanding the biological sequelae of childhood adversity––and how those sequelae differ depending on sensitive periods in exposure––represents the first step towards the development of targeted strategies designed to disrupt the processes linking adversity to psychiatric diseases as early in life course as possible.

## Acknowledgments

This research was specifically funded by the National Institute of Mental Health of the National Institutes of Health under Award Numbers K01MH102403 and 1R01MH113930 (Dunn) and under R03AG051877 and 3R03AG051877-02S1 (Wolf). The content is solely the responsibility of the authors and does not necessarily represent the official views of the National Institutes of Health, the U.S. Department of Veterans Affairs, or the United States Government. The authors thank Alice Renaud for her assistance in preparing this manuscript for publication. We are extremely grateful to all the families who took part in this study, the midwives for their help in recruiting them, and the whole ALSPAC team, which includes interviewers, computer and laboratory technicians, clerical workers, research scientists, volunteers, managers, receptionists and nurses. The UK Medical Research Council and the Wellcome Trust (Grant ref: 102215/2/13/2) and the University of Bristol provide core support for ALSPAC. This publication is the work of the authors who will serve as guarantors for the contents of this paper. A comprehensive list of grants funding is available on the ALSPAC website (http://www.bristol.ac.uk/alspac/external/documents/grant-acknowledgements.pdf).

